# Amino Sugars Reshape Interactions between *Streptococcus mutans* and *Streptococcus gordonii*

**DOI:** 10.1101/2020.06.22.166272

**Authors:** Lulu Chen, Alejandro R. Walker, Robert A. Burne, Lin Zeng

## Abstract

Amino sugars, particularly glucosamine (GlcN) and N-acetylglucosamine (GlcNAc), are abundant carbon and nitrogen sources that are continually supplied in host secretions and in the diet to the biofilms colonizing the human oral cavity. Evidence is emerging that these amino sugars provide ecological advantages to beneficial commensals over oral pathogens and pathobionts. Here, we performed transcriptome analysis on *Streptococcus mutans* and *Streptococcus gordonii* growing in single-species or dual-species cultures with glucose, GlcN or GlcNAc as the primary carbohydrate source. Compared to glucose, GlcN caused drastic transcriptomic shifts in each bacterium when they were cultured alone. Likewise, co-cultivation in the presence of GlcN yielded transcriptomic profiles that were dramatically different than the single-species results from GlcN-grown cells. In contrast, GlcNAc elicited only minor changes in the transcriptome of either organism in single- and dual-species cultures. Interestingly, genes involved in pyruvate metabolism were among the most significantly affected by GlcN in both species, and these changes were consistent with measurements of pyruvate in culture supernates. Differing from a previous report, growth of *S. mutans* alone with GlcN inhibited expression of multiple operons required for mutacin production. Co-cultivation with *S. gordonii* consistently increased the expression by *S. mutans* of two manganese transporter operons (*slo* and *mntH*) and decreased expression of mutacin genes. Conversely, *S. gordonii* appeared to be less affected by the presence of *S. mutans*, but did show increases in genes for biosynthetic processes in the co-cultures. In conclusion, amino sugars profoundly alter the interactions between a pathogenic and commensal streptococcus by reprogramming central metabolism.

**IMPORTANCE:** Carbohydrate metabolism is central to the development of dental caries. A variety of sugars available to dental microorganisms influence the development of caries by affecting the physiology, ecology, and pathogenic potential of tooth biofilms. Using two well-characterized oral bacteria, one pathogen (*Streptococcus mutans*) and one commensal (*Streptococcus gordonii*) in a RNA deep-sequencing analysis, we studied the impact of two abundant amino sugars on bacterial gene expression and interspecies interactions. The results indicated large-scale remodeling of gene expression induced by GlcN in particular, affecting bacterial energy generation, acid production, protein synthesis, and release of antimicrobial molecules. Our study provides novel insights into how amino sugars modify bacterial behavior, information that will be valuable in the design of new technologies to detect and prevent oral infectious diseases.

## INTRODUCTION

Amino sugars are particularly abundant in the human oral cavity. In particular, N-acetylglucosamine (GlcNAc) and glucosamine (GlcN) decorate host proteins, such as mucins and basic proline-rich glycoproteins (PRG), and are the fundamental building blocks of the cell envelopes of bacteria and fungi (1, 2). These carbohydrates are also common in foodstuffs consumed by humans. GlcN and GlcNAc can be released from polymers through the collective activities of microbial glycohydrolases, providing a steady supply of carbon- and nitrogen-sources to the oral microflora, especially while the host is fasting. Most known oral bacteria have the capacity, or carry the necessary genes, for metabolizing exogenous sources of amino sugars. GlcN and GlcNAc catabolism generally requires a specialized membrane transporter, often a permease of the sugar: phosphotransferase system (PTS); a GlcNAc-6-P deacetylase (NagA) that converts GlcNAc-6-P into GlcN-6-P; and a GlcN-6-P deaminase (NagB) that converts GlcN-6-P into fructose-6-P (F-6-P). F-6-P can enter the Embden–Meyerhof–Parnas (EMP) pathway, while the molecule of ammonia released from GlcN or GlcNAc can complex with a proton inside or outside the cell to raise the pH, or can be funneled into amino acid biosynthesis. Metabolism of GlcNAc also releases a molecule of acetate, which can enter catabolic or anabolic processes, depending on the organism and its bioenergetic and nutritional needs (3).

With only a partial tricarboxylic acid (TCA) cycle, the dental caries pathogen *Streptococcus mutans* depends almost exclusively on the EMP pathway for production of ATP (4, 5). Under carbohydrate-excess and low-oxygen conditions, the bacterium converts the product of glycolysis, pyruvate, primarily into lactate, via a lactate dehydrogenase (LDH) complex, with concomitant conversion of NADH to NAD^+^ (6). As a fairly strong organic acid, lactate drives down the environmental pH, accelerating the initiation and progression of human dental caries. LDH in *S. mutans* is allosterically activated by the glycolytic intermediate fructose-1,6-bisphosphate (F-1,6,-bP) and functions optimally in a pH range of 5.5 to 6.3 for pyruvate reduction (6, 7). Under carbohydrate-limiting conditions, metabolism of pyruvate proceeds mainly by pyruvate dehydrogenase (PDH) or by an oxygen-sensitive pyruvate formate lyase (PFL); yielding acetyl-CoA and converting NAD^+^ into NADH (8, 9). Many commensal oral streptococci, e.g., *Streptococcus gordonii* and *Streptococcus sanguinis*, express a pyruvate oxidase enzyme (SpxB) that converts pyruvate into acetyl-phosphate (AcP) with the concomitant production of hydrogen peroxide (H_2_O_2_) (10). *S. mutans* is known to be sensitive to H_2_O_2_, with inhibition of certain enzymes, cellular processes and growth being evident at the levels of H_2_O_2_ that can be produced and well-tolerated by many commensal streptococci. Lacking SpxB, *S. mutans* converts acetyl-CoA into AcP using the enzyme phosphotransacetylase (Pta) (11). AcP can be acted on by an acetate kinase (AckA) in the presence of ADP to produce acetate and ATP. Notably, many bacteria allow the reverse reaction, converting acetate into AcP and eventually into acetyl-CoA. In addition to its involvement with pyruvate and acetyl-CoA metabolism, AcP appears to have important roles in gene regulation and can serve as a substrate for certain two-component signal transduction systems (12).

GlcN and GlcNAc can significantly impact bacterial fitness and interbacterial interactions in streptococci that are abundant in dental biofilms (13, 14). For example, amino sugars enhanced the ability of two health-associated commensal bacteria, *S. gordonii* and *S. sanguinis*, to outcompete *S. mutans in vitro* in planktonic cultures and in biofilms (13, 14). These observations were generally attributed to: i) more efficient utilization of amino sugars, GlcNAc in particular, by the commensals; ii) increased release of H_2_O_2_ by the commensals growing on amino sugars; iii) and moderation of acidification of the environment by release of ammonia from the amino sugars and through increased expression of the arginine deiminase (AD) system of the commensals growing on amino sugars (15). Conversely, growth on amino sugars was also reported to result in enhanced acid tolerance and increased bacteriocin (mutacin) production by *S. mutans* (13, 16). Furthermore, in a human saliva-derived *ex vivo* microcosm model, GlcNAc but not GlcN was shown to inhibit the persistence of *S. mutans* (13). In light of the profound impact of amino sugars on both beneficial properties of commensals and virulence-related attributes of *S. mutans*, a more comprehensive understanding of the effects of amino sugars on these oral bacteria is needed, particularly in the context of the ecological dynamics that influence the balance between oral health and diseases. By applying RNA deep sequencing (RNA-Seq) to *S. mutans* UA159 and *S. gordonii* DL1 in a batch-culture model, we uncovered molecular mechanisms underlying physiological changes and behaviors by these bacteria that are unique to amino sugar effects and to co-cultivation when amino sugars are present. The study reveals significant physiological reprogramming in both bacteria that can guide future work to manipulate bacterial ecology to control oral infectious diseases.

## RESULTS AND DISCUSSION

### RNA-Seq sample clustering

The two main objectives of this study were to catalogue the impacts of GlcN and GlcNAc, as compared to glucose, on the transcriptomes of *S. mutans* UA159 and *S. gordonii* DL1, and to understand how these carbohydrates affect competition between these bacteria. To achieve these goals, each organism was grown in TV base medium formulated with one of the three carbohydrates, either in single- or dual-species cultures. For each condition, there were three biological replicates. TV medium was chosen over synthetic FMC (17) and TY medium for specific reasons: *S. mutans* experiences significant delays in growth on FMC-GlcNAc when transferred from a glucose-based medium (14), and TV lacks the yeast extract in TY that adds contaminating carbohydrates that could influence the interpretation of the results. *In toto*, 27 mRNA-enriched samples and cDNA libraries were prepared, 18 from single-species cultures and 9 from co-cultures. After cDNA sequencing and data analysis, it was determined that there was a sufficient quantity of high-quality reads and sufficient coverage of the genomes of both organisms under all conditions tested, with each individual library yielding an average of about 9 million reads. After mapping all qualified reads to annotated genes in both genomes, 18 RNA-Seq samples of each bacterium were subjected to correlation, clustering, and principal component analyses (PCA) to assess their overall sample quality and relationships (Fig. 1 and Fig. S1). As visualized in Fig. 1, the replicates from each group showed good consistency in gene expression patterns. Interestingly, the single-species cultures grown on GlcN showed the greatest divergence from the groups grown on glucose or GlcNAc. Co-cultivation of *S. mutans* and *S. gordonii* with GlcN also showed the greatest impact on both transcriptomes relative to the single-species cultures on GlcN. In contrast, each organism expressed more similar transcriptomes when growing on glucose or GlcNAc as single-species or in co-cultures. The tendency for GlcN cultures to diverge from all other samples was even better illustrated by the results of the principal component analysis (PCA) in Fig. 1C,D.

**Fig. 1.**
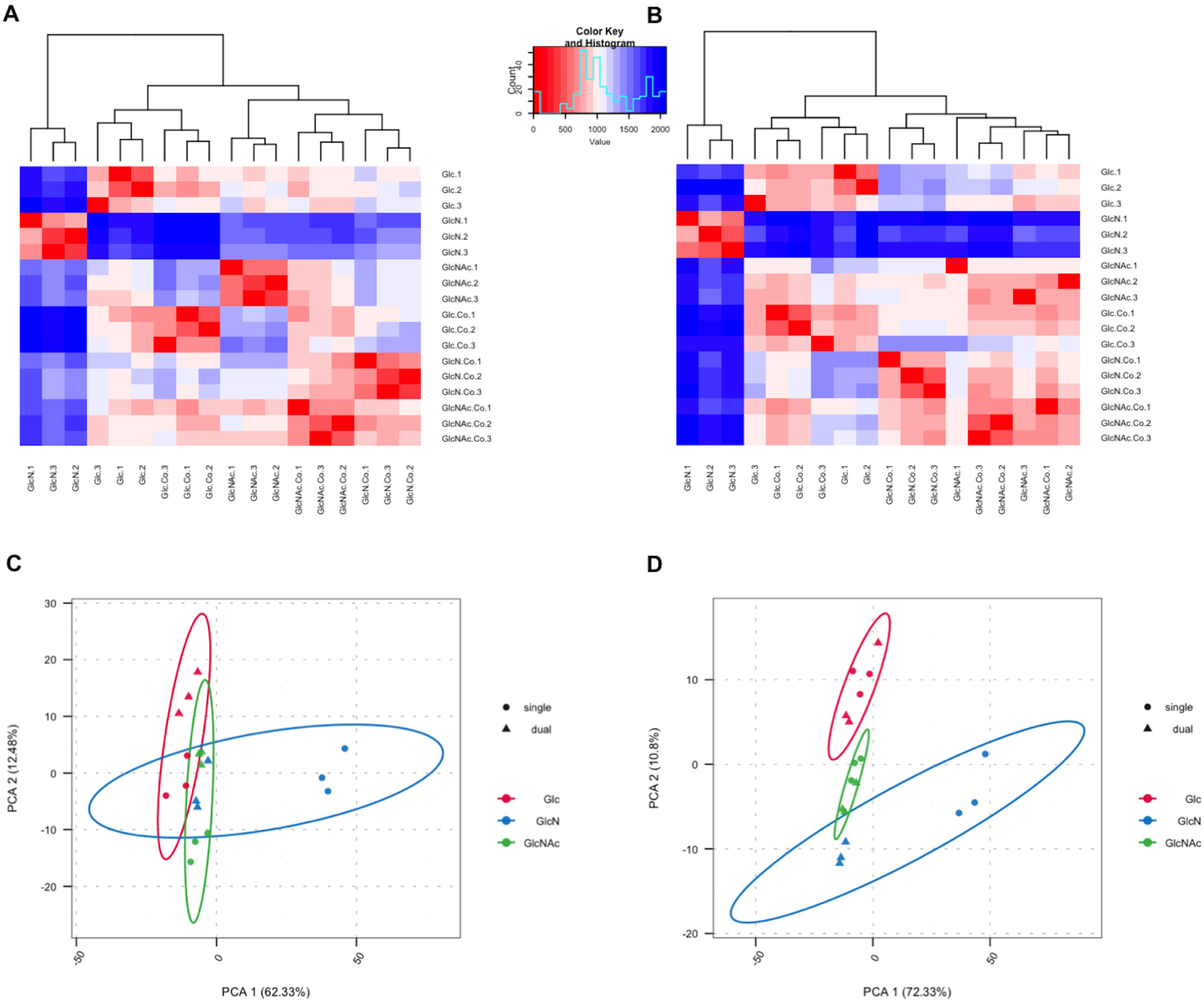
Clustering (A, B) and principal component analysis (C, D) of 18 RNA-Seq samples for *S. mutans* (A, C) and *S. gordonii* (B, D). (A, B) Heatmap of the cluster analysis shows the Euclidean distances between the samples as calculated from the variance-stabilizing transformation of the count data. (C, D) The circles in PCA diagrams represent single-species samples and the triangles represent mixed-species samples. The red, blue, and green symbols represent samples grown on glucose (Glc), GlcN, and GlcNAc, respectively.

### Differential expression analysis

With carbohydrates and cultivation conditions as the two independent variables, we carried out 5 pair-wise comparisons for each bacterium. For single-species samples, GlcN and GlcNAc conditions were each contrasted against the glucose (Glc) samples; for co-cultures, results from each carbohydrate were compared with results from single-species cultures grown on the same carbohydrate. The cutoff for identifying differentially expressed genes (DEG) was a false discovery rate (FDR) <0.01 and a |Log_2_(fold change)| ≥1.5 (or fold-change ≥2.8) in the number of reads for a given mRNA. These results are presented in volcano plots and Venn diagrams (Fig. 2&3) with the lists of DEG included in supplemental materials (Tables S1 through S7). To better understand the biological processes associated with the DEG, functional enrichment analysis of Gene Ontology (GO) was performed using DEG for identification of specific pathways, which are shown as KEGG maps (see supplemental materials). Subsequently, a subset of genes from both bacteria were selected for validation using RT-qPCR (Fig. 4).

**Fig. 2.**
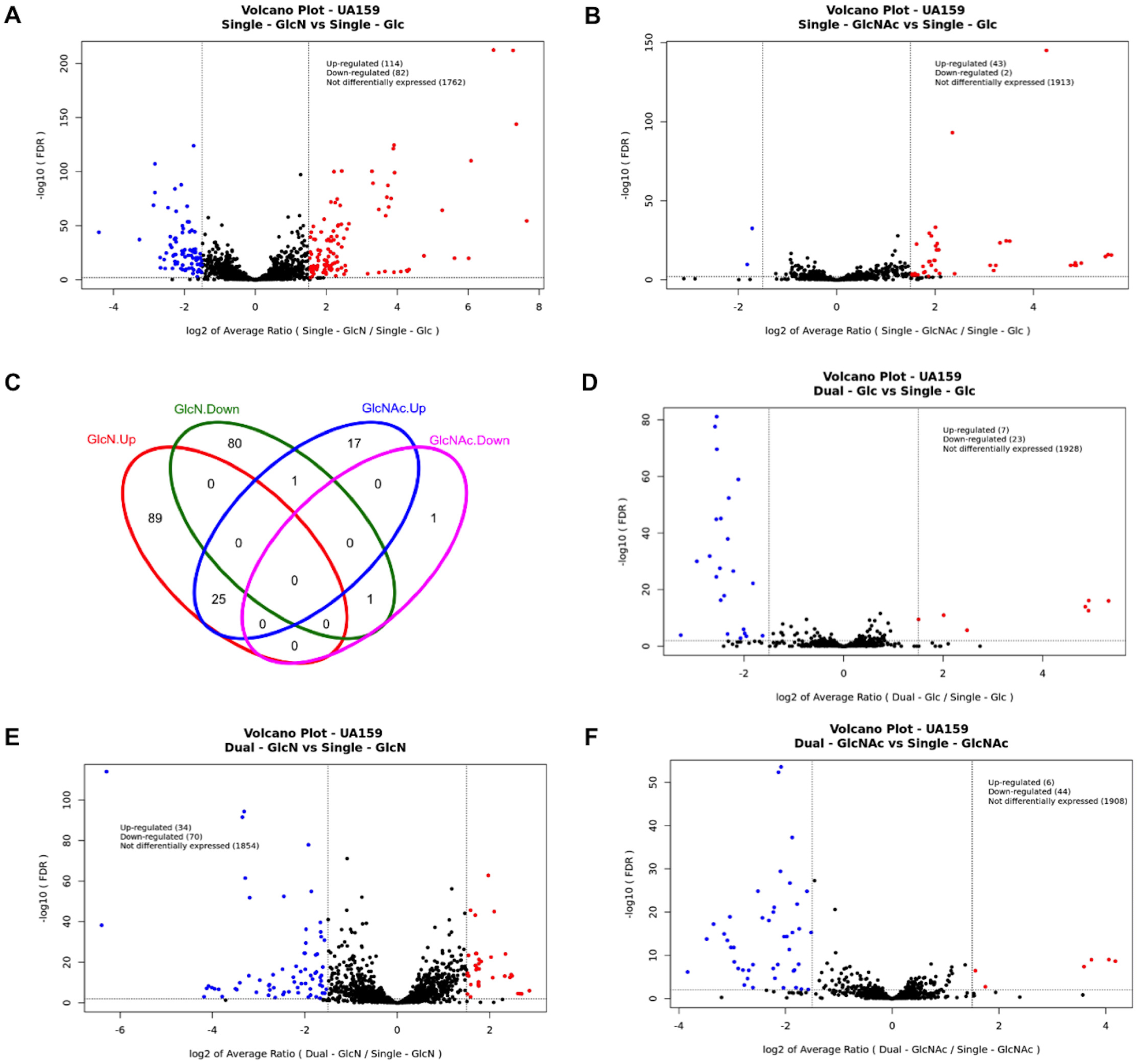
Differentially expressed genes (DEG) from *S. mutans* UA159 identified by RNA-Seq. Qualified reads were mapped to the genome of UA159, counted for each annotated gene, and normalized across the entire transcriptome. The DEG were identified by comparison of single-species cultures between GlcN and Glc (A), or GlcNAc and Glc (B), and by comparison of mixed-species cultures against single-species cultures, each grown on Glc (D), GlcN (E), or GlcNAc (F). Red circles depict DEG with increased expression and blue circles depict those with decreased expression. All DEG from panels A and B are presented as one Venn diagram (C).

**Fig. 3.**
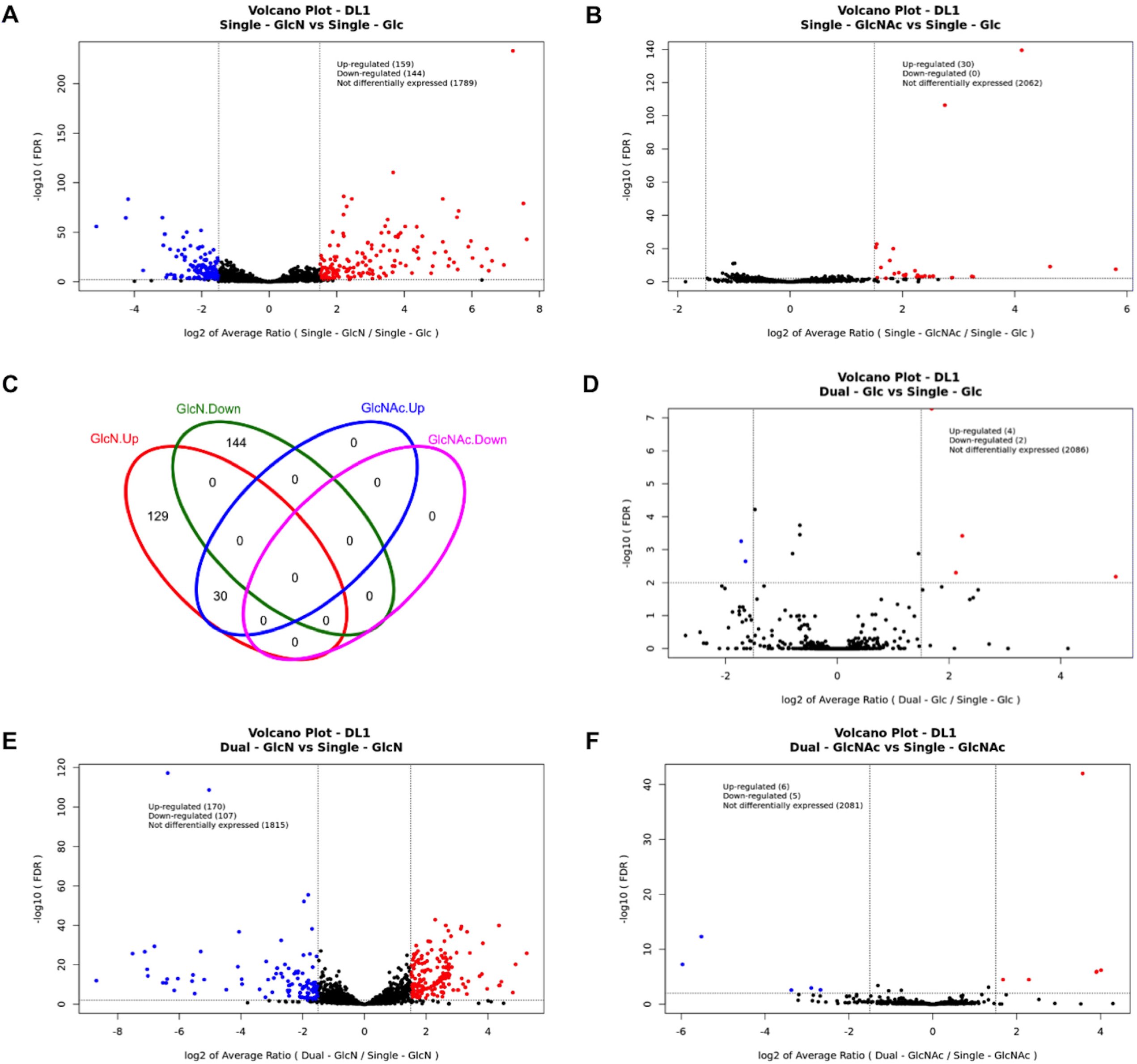
DEG from *S. gordonii* DL1 identified by RNA-Seq. The DEG were identified by comparison of single-species cultures between GlcN and Glc (A), or GlcNAc and Glc (B), and by comparison of mixed-species cultures against single-species cultures, each grown on Glc (D), GlcN (E), or GlcNAc (F). Red circles depict DEG with increased expression and blue circles depict those with decreased expression. All DEG from single-species cultures are presented as one Venn diagram (C).

**Fig. 4.**
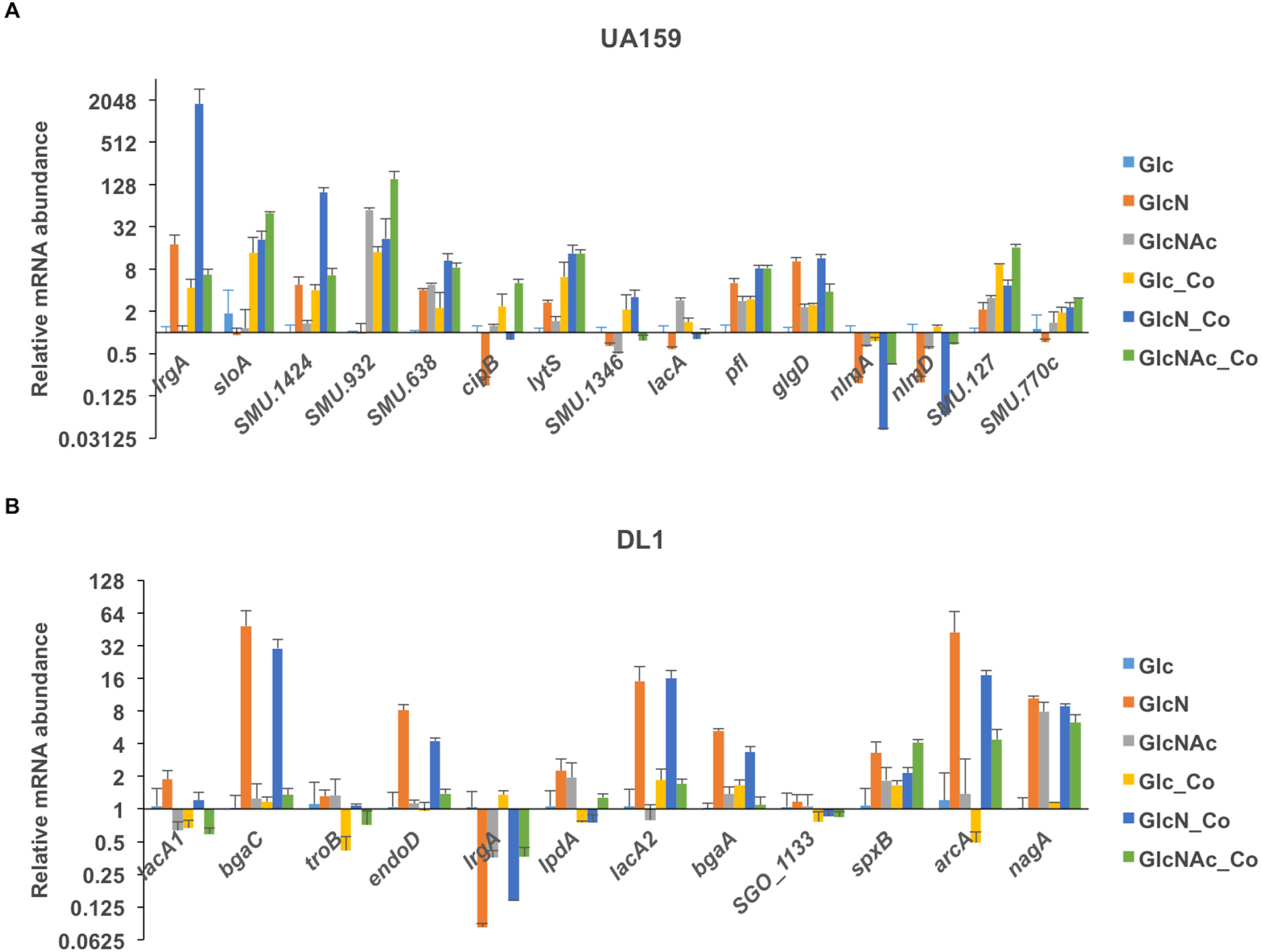
RT-qPCR of differentially expressed genes from *S. mutans* (A) and *S. gordonii* (B). The results of each gene, including single-species and co-cultures (-Co), are presented as relative mRNA abundance by comparison to levels measured in single-species cultures supported by glucose (Glc). Each bar represents the average of three biological repeats, with the error bar denoting the standard deviation. Culture conditions for RNA isolation differed for the RNA-Seq and RT-qPCR as detailed in the methods and results sections.

#### i. *S. mutans* in single-species cultures

Growth of *S. mutans* on GlcN and GlcNAc can elicit some similar behaviors. In particular, both sugars enhanced acid tolerance of *S. mutans*, presumably due to cytoplasmic alkalization by release of ammonia (16), along with increased mutacin production, as assessed in a deferred antagonism assay on TY agar formulated with the different sugars (13). Conversely, there are GlcNAc-specific effects that include an extended lag phase when *S. mutans* is transferred from glucose-containing media to a medium with GlcNAc as the primary carbohydrate source. (14). There is also a significant reduction in persistence of the bacterium in a human-derived *ex vivo* consortium biofilm model provided with GlcNAc as the main carbohydrate (13).

##### UA159 on GlcN alone

When *S. mutans* UA159 was cultivated alone in TV-GlcN as compared to TV-glucose, 114 annotated genes showed significantly increased expression while 82 others had decreased expression (Fig. 2A, Fig. 4A, and Table S1 for a full list of DEG). The differentially expressed genes mainly included pathways for carbohydrate metabolism: PTS, glycolytic enzymes, pyruvate metabolism, and gene products for use of cellobiose, galactose, lactose, sucrose, starch and glycogen. Additionally, amino acid transport, biosynthesis and metabolism of lysine, serine, threonine and methionine, along with nucleotide metabolism and bacteriocin production were substantially affected by growth in GlcN, compared with cells growing on glucose (Fig. S2 for KEGG visualization). Notably, two operons in *S. mutans* UA159 showed the greatest increase in mRNA levels when growing on GlcN: the pyruvate dehydrogenase operon (PDH, SMU.1421~SMU.1425; increased by 68~147-fold), and a pyruvate transporter operon (*lrgAB*, SMU.574c~SMU.575c; increased by 137~156-fold)(18) together with the genes encoding the two-component signal transduction system that activates *lrgAB* (*lytTS*, SMU.576~SMU.577; increased by 5-fold)(19). It was recently discovered that *S. mutans* excretes pyruvate as an overflow metabolite under glucose-rich conditions, only to reinternalize the pyruvate once glucose is depleted in the environment (18). Another apparent orthologue of the PDH complex, SMU.127-SMU.130, also showed about a 2-fold increase in mRNA levels (Fig. 4A). Genes homologous to this particular PDH operon are widely conserved in multiple streptococcal species, including *S. gordonii*, *Streptococcus pneumoniae*, and group A *Streptococcus*. Two other genes encoding enzymes for pyruvate metabolism, pyruvate formate lyase (*pfl*, SMU.402) and PFL-activating enzyme (*pflC*, SMU.490)(9) were upregulated about 4-fold in cells grown on GlcN, compared to cells grown on glucose. Under carbohydrate-rich and low-oxygen tension conditions, *S. mutans* generally depends on glycolysis and lactate generation to meet its demand for energy production and growth. These alternate pyruvate metabolic pathways are often activated when energy sources are limiting, as use of pyruvate by these routes yields greater amounts of ATP per molecule of carbohydrate catabolized. Also of significance, a number of genes encoding for mutacins and related proteins, including *nlmA*, *nlmB*, SMU.152-SMU.153, *nlmC/cipB*, *nlmD*, *nlmE*, and *nlmT* showed decreased mRNA levels in GlcN-grown cultures (Fig. 4A and Table S1). This was a surprising finding as we reported previously that *S. mutans* UA159 showed enhanced killing of an indicator *S. sanguinis* SK150 in a deferred antagonism assay using TY-agar, not the TV medium used here, when supplemented with amino sugars (13). Insights into why these differences exist are provided below.

##### UA159 on GlcNAc alone

When UA159 was cultured on TV-GlcNAc, 43 genes showed higher mRNA levels and only 2 genes showed lower mRNA levels, relative to cultures on glucose (Fig. 2B, Fig. 4A, and Table S2). Among these genes with the highest increase in expression were an ABC-transporter operon (SMU.651c~SMU.653c; >40-fold increase) with unknown substrate(s), along with a putative amino acid transporter operon (SMU.932-SMU.936; 23-28-fold increase) and its nearby genes (SMU.930c, 8-fold increase; SMU.937-SMU.939, ~2-fold increase). A significant portion of these 43 genes were involved in energy metabolism, followed by genes annotated for nucleotide metabolism; most notably a 20-gene cluster, ranging from SMU.29 to SMU.59 and annotated for purine biosynthesis, showed 2 to 6-fold increases in mRNA levels.

There were 25 genes that were more highly expressed in both GlcN and GlcNAc (Fig. 2C, Table S3) compared to glucose. Most notable were the *nagA* and *nagB* genes encoding for GlcNAc-6-P deacetylase and GlcN-6-P deaminase, respectively. The expression of these genes is known to be induced by growth on these amino sugars (16). Some genes linked to *nagAB* were also differentially expressed. Other genes showing increased expression under both conditions were the tagatose pathway (SMU.1492-SMU.1496) and the aforementioned two ABC transporter operons (SMU.932-SMU.936 and SMU.651c-SMU.653c). As reported previously, growth on amino sugars only down-regulated the *glmS* gene, encoding for the GlcN-6-P synthase (16). In light of the results showing reduced expression of mutacin genes on GlcN, we further examined the expression of these genes in GlcNAc-grown cultures. As shown in Fig. 4A, compared to glucose, GlcNAc reduced the message levels of *nlmAB* and *nlmD*, but the reduction was not as significant as that seen with growth on GlcN. No effect of GlcNAc was noted for *nlmC/cipB*.

Metabolism of GlcN differs from that of glucose primarily in the deamination of GlcN-6-P by NagB, thereby releasing an ammonia molecule intracellularly. In addition to generation of ammonia, degradation of GlcNAc by *S. mutans* also requires deacetylation of GlcNAc-6-P, yielding a molecule of acetate. Since expression of both *nagA* and *nagB* is co-regulated by the GntR-type regulator NagR in association with the allosteric effector GlcN-6-P produced by catabolism of both amino sugars (20), the drastically disparate gene expression patterns among these three carbohydrate conditions are most likely the outcome of cellular responses to distinct physiological conditions. Under these batch culture conditions, where carbohydrate is not limiting, lactate dehydrogenase LDH is likely the main pathway for metabolizing pyruvate generated by glycolysis, concurrently regenerating NAD^+^ and yielding lactate as the end product. The enzymatic activity of LDH in *S. mutans* functions optimally between pH 5.5 and 6.3, and the cells also have more LDH protein when growing at pH 5.5 than 7.0 (7). We reason that the release of ammonia by the degradation of GlcN and GlcNAc increases intracellular pH (16) and reduces LDH activities, which in turn raises intracellular pyruvate levels, redirecting carbon flow from generation of lactate to the PDH and/or PFL pathways for generation of acetate or acetate and formate, respectively. Since metabolism of GlcNAc also releases a molecule of acetate, acetate accumulation could create negative feedback pressure upstream of PDH and/or PFL (11), and thus the forward reactions of these alternative pyruvate dissimilation pathways may be inhibited; possibly allowing accumulation of pyruvate and even acetyl phosphate. It is also worth noting that *S. mutans* growing on GlcN generated more ammonia and reached higher final pH values than those growing on GlcNAc (16). This difference could be due to the faster catabolism of GlcN or more ammonium ions being directed into anabolic pathways in cells growing on GlcNAc. Another perhaps equally important outcome of reduced LDH activity could arise from changes in the NAD^+^/NADH ratios. Balancing NAD^+^/NADH is critical in *S. mutans*, and imbalances could disrupt glycolysis and intracellular redox balance, either of which could have a profound impact on global gene expression (21). One example of this likely compensatory response is increased expression of the glucose-PTS (EII^Man^, *manLMN*) that was shown to be mediated by the redox sensor Rex (22). Further examination of the RNA-Seq data indicated that the *ldh* gene in *S. mutans* showed a modest reduction (~40%) in expression in cells growing on GlcN, as compared to glucose, whereas the *pfl*, *pflC* and the second PDH operon (SMU.127-SMU.130) showed significantly increased expression in cells growing on GlcNAc: 3-fold, 3-fold, and 2-fold, respectively (Fig. 4A and RNA-Seq data not shown). Expression of the *manLMN* operon also was increased by about 2-fold in both GlcN and GlcNAc cultures. However transcripts of the PDH operon (SMU.1421-SMU.1425) showed no change in GlcNAc.

To further test the hypothesis that amino sugars impacted pyruvate metabolism, we measured the extracellular pyruvate levels in *S. mutans* UA159 cultured on glucose, GlcN or GlcNAc in different growth phases. As pyruvate is known to be excreted into the environment by *S. mutans*, we expected that a reduction in LDH activity could increase the pyruvate concentrations in the cells and culture supernates. The results (Fig. 5ABC) showed that as the optical density of the cultures increased, so did the levels of extracellular pyruvate, regardless of the carbohydrate source. As the glucose cultures entered stationary phase, the pyruvate levels declined rapidly and all but disappeared in a matter of hours; likely due to reinternalization by LrgAB transporter (18). In comparison, the peak extracellular pyruvate levels in cultures growing on GlcN were about 33% higher than in glucose cultures (*p* <0.001). Furthermore, although it took slightly longer to reach the peak levels on GlcN, the rate at which the pyruvate levels declined afterwards was also significantly slower. In GlcNAc-supported cultures, the peak pyruvate levels were again higher (by 30~40%) than glucose cultures (*p* <0.05), however the decline followed a rapid trajectory, similar to cells growing on glucose.

**Fig. 5.**
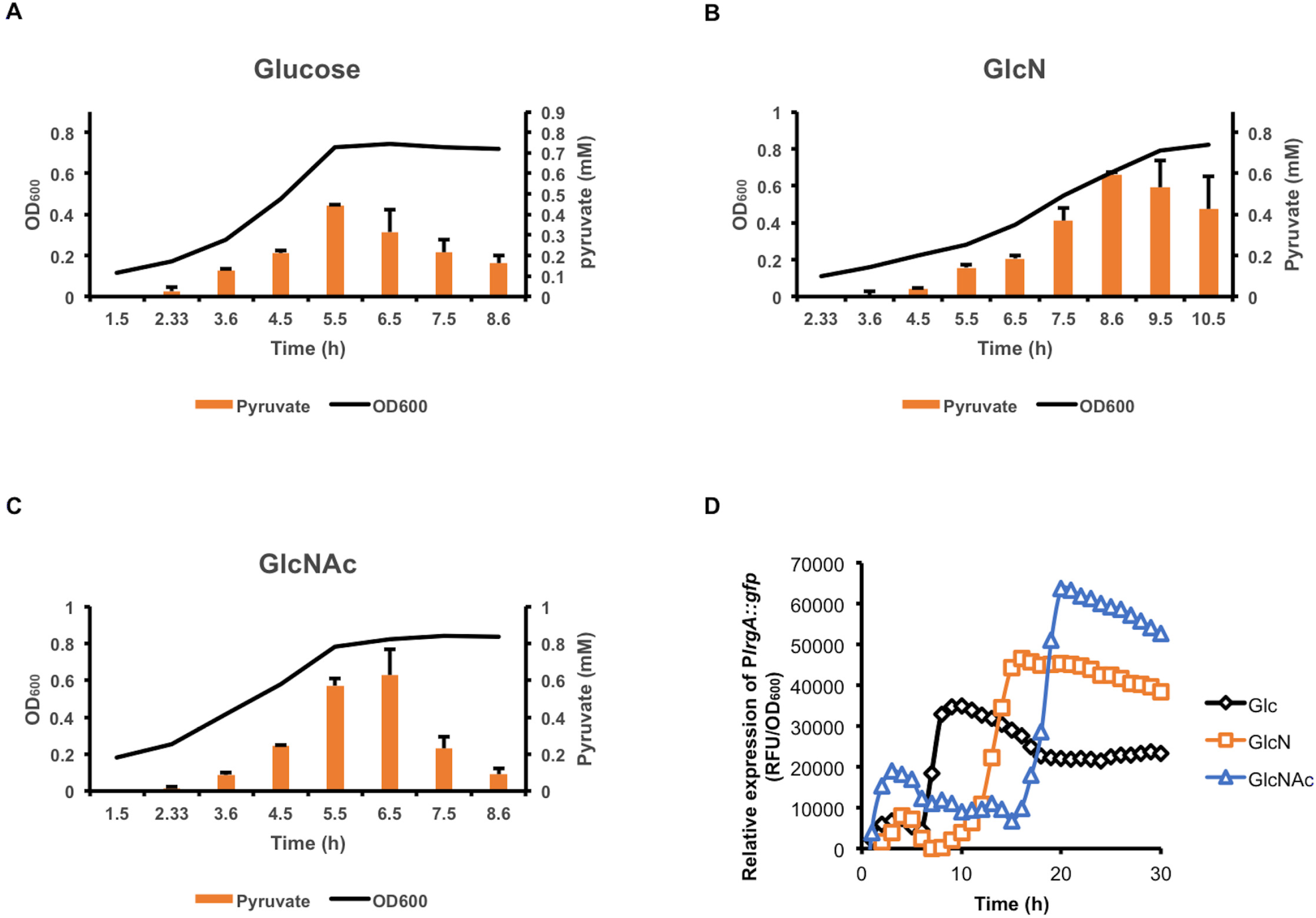
Measurements of extracellular pyruvate levels and P*lrgA∷gfp* expression in *S. mutans* cultures. *S*train UA159 was cultured in an FMC base medium supplemented with 10 mM glucose (A), GlcN (B), or GlcNAc (C), with pyruvate levels measured at regular intervals at various stages of growth. The optical density (OD_600_, black line) was plotted along with the corresponding pyruvate levels (orange bars) against the time. Each bar represents the average of three biological repeats, with the error bar denoting the standard deviation. (D) UA159/P*lrgA∷gfp* was similarly cultured on FMC and monitored for growth and GFP signals for a period of 30 h.

The transcription of the *lrgAB* genes has been shown to be induced by the release of pyruvate (18). As such, we performed a promoter∷reporter fusion analysis on a UA159 derivative carrying a P*lrgA∷gfp* fusion (18) by monitoring GFP levels in cells growing on glucose, GlcN or GlcNAc. Our results (Fig. 5D) showed increased *lrgA* promoter activities when cells were growing on GlcN and especially on GlcNAc, as compared to glucose, although the peak promoter activities appeared later in GlcN and even later in GlcNAc. Together, these results are consistent with our hypothesis that reprograming of pyruvate metabolism likely occurred as a result of the differences in physiological conditions in cells catabolizing the three different carbohydrates.

#### ii. *S. gordonii* cultured alone

Previous research on amino sugar metabolism by *S. gordonii* and related commensals (13, 14) has shown enhanced competitiveness of the commensals against the oral pathogen *S. mutans*, primarily through increased production of H_2_O_2_ (with both amino sugars) and higher expression of the arginine deiminase pathway, mostly notably with GlcN. However, little is known regarding the underlying mechanisms responsible for these important phenotypes or the more global responses of commensals to these sugars.

##### DL1 on GlcN alone

When single-species cultures of *S. gordonii* DL1 were growing on TV-GlcN, 159 genes showed increased expression and 144 showed decreased expression, compared to cells growing in TV-glucose (Fig. 3A, Fig. 4B and Table S4). The majority of these DEGs belonged to carbohydrate metabolic pathways, but significant proportions also involved nucleotide and amino acid metabolic networks (Fig. S3 for KEGG visualization). Similar to *S. mutans* growing on GlcN, most of these carbohydrate metabolic genes showed significantly increased expression in the presence of GlcN, including: two extracellular β-galactosidase, *bgaC*/EII operon (SGO_0043~SGO_0048, increased by 29~210-fold)(23, 24) and *bgaA* (SGO_1486, increased by 9-fold)(24), two putative glucan-binding proteins (SGO_0477~SGO_0478, increased by ~40-fold), two Leloir pathway genes (*galKT*, increased by 6~7-fold), an endo-β-N-acetylglucosaminidase D (*endoD*, 6-fold increase)(23), a 17-gene, two-cluster system encoding for the tagatose pathway and PTS permeases required for metabolism of lactose and galactose (SGO_1512-SGO_1526, increase ranging from 5 to 100-fold)(24), a putative glycogen biosynthetic operon (SGO_ 1549-SGO_1554, increased by 3-5-fold), a trehalose metabolic operon (SGO_1652-SGO_1653, increased by 69- and 116-fold, respectively), a putative fructose/mannose PTS operon (SGO_1890-SGO_1893, increased by ~3-fold), and a putative sugar-binding ABC transporter operon (SGO_ 1763-SGO_1765, increased by 22-28-fold). Also increased in expression were two enzymes responsible for producing H_2_O_2_: the pyruvate oxidase *spxB* cluster (SGO_0289~SGO_0293, increased by 5~20-fold) and the NADH oxidase gene (*nox*, SGO_1167) showed 5-fold increases. The only pyruvate dehydrogenase operon in *S. gordonii* (SGO_1130-SGO_1133; SGO_1130 = *lpdA*, Fig. 4B) showed 4- to 7-fold increases in expression. However, the expression of two *cidAB*/*lrgAB* orthologue genes (SGO_1268~-SGO_1269) decreased by ~20-fold. Pyruvate overflow metabolism has been reported in *S. gordonii* (25), although it is not known if the *cidAB*/*lrgAB* gene products contribute to pyruvate metabolism in ways similar to that of *S. mutans*.

Among DEGs related to amino acid metabolism were the arginine deiminase (AD) operon (*arcABCDT*), which showed a 3- to 6-fold increase in expression in GlcN, compared to glucose. This result is consistent with the reported increase in AD activities of *S. gordonii* growing on amino sugars (14). It is also known that *arc* operon expression in DL-1 is strongly induced by arginine, repressed by oxygen and sensitive to catabolite repression (26). Conversely many genes required for protein biosynthesis, most notably 17 ribosomal proteins, 18 tRNAs, and 5 amino acid-tRNA synthetases, showed significantly reduced expression (Table S4). Significant number of genes from the nucleotide metabolism network also showed reduced expression in GlcN-grown cells. The simplest interpretation of these findings is that *S. gordonii* growing on GlcN scaled down the biosynthesis of proteins and nucleotides, which are required for growth, while directing more carbon flux to energy production. Similar to *S. mutans*, it is probably safe to assume that metabolism of GlcN, as well as GlcNAc, increases the intracellular pH of *S. gordonii*. Although the effects of pH on LDH from *S. gordonii* is not well characterized, LDH activity is likely lower when intracellular pH is elevated (6). This notion is supported by the increased expression of pyruvate oxidizing enzymes PDH and SpxB, and increased H_2_O_2_ production in the presence of GlcN (Table S4 and Fig. 4B)(13).

##### DL1 on GlcNAc alone

When *S. gordonii* DL1 was cultured on TV-GlcNAc, there were only 30 genes that showed increased expression in comparison to glucose cultures, and no genes were identified that had decreased expression (Fig. 3B, Fig. 4B and Table S4). Interestingly, all 30 genes also showed increased expression in GlcN cultures (Fig. 3C). The 30 genes are almost exclusively dedicated to carbohydrate metabolism, encompassing the *nag* pathway, the lactose/galactose cluster, the trahalose operon, the extracellular β-galactosidases BgaA/BgaC, and the endo-β-N-acetylglucosaminidase EndoD. For comparison, in *S. mutans* these two amino sugar regulons overlap by 25 genes, yet the GlcNAc regulon of *S. mutans* contained an additional 18 unique DEGs that were not DEGs in cells growing on GlcN.

Current research supports roles for EndoD and BgaC in the ability of *S. gordonii* to grow on saliva by sequentially releasing monosaccharides from the basic proline-rich glycoproteins, in concert with the activity of the β-*N*-acetylglucosaminidase StrH (23). Since amino sugars and galactose are likely released during glycoprotein degradation, but GlcN and GlcNAc were the only carbohydrates used in our cultures, it is possible that amino sugars or their metabolic derivatives are sufficient to activate the transcription of these secreted glycohydrolases, along with the tagatose pathway that is required for catabolizing galactose. Alternatively, some of these genes showing altered expression on both amino sugars may be due to shifts in pyruvate metabolism, as discussed above.

#### iii. *S. mutans* and *S. gordonii* cultured together

##### Co-culture on Glc

Co-cultivation with *S. gordonii* on glucose presented a clear oxidative stress to *S. mutans*, apparently due to H_2_O_2_ production, as UA159 showed enhanced expression in the *slo* operon (SloABCR, SMU.182-SMU.186), and a two-gene cluster, SMU.768c-SMU.770c that encodes for a metal ion transporter MntH (Fig. 4A, Fig. 6, and Table S5), in comparison to *S. mutans* growing in monoculture on glucose. SloR is a manganese-responsive metalloregulator that is required for *S. mutans* response to oxidative and acidic stressors (27, 28), and SloABC is a Mn^2+^/Fe^2+^ transporter that works in concert with MntH as the primary means of Mn^2+^ uptake in *S. mutans* (29, 30). The capacity of SloR to regulate both the *slo* operon and *mntH*, along with the known positive effects of Mn^2+^ on *S. mutans* aero-tolerance, suggest that H_2_O_2_ production by DL1 may have triggered the induction of these gene. At the same time, co-cultivation with *S. gordonii* reduced expression of various mutacin genes, namely the *nlmAB* operon, *nlmD*, and the cluster SMU.1902c-SMU.1914c that includes *nlmC*/*cipB*. The decrease in mutacin gene expression is likely, at least in part, due to production by DL-1 of the protease challisin (Sgc). Sgc is secreted by DL1 and can degrade competence-stimulating peptide (CSP), which is required for mutacin expression (31). Notably, the pyruvate transporter gene *lrgA* was also identified as being down-regulated in co-cultured *S. mutans*.

**Fig. 6.**
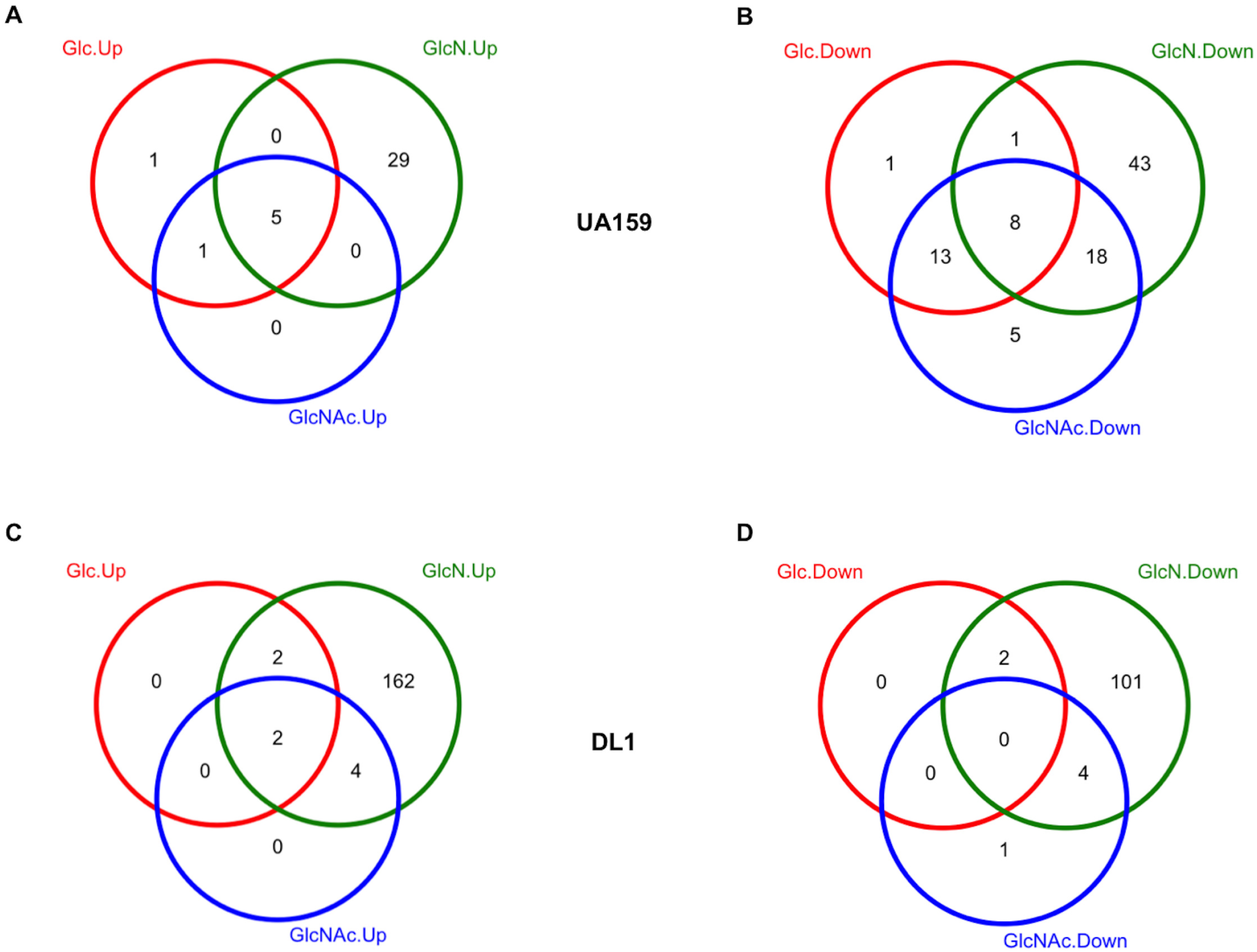
Venn diagrams comparing DEG identified in *S. mutans* UA159 (A, B) and *S. gordonii* DL1 (C, D) in mixed-species cultures in comparison to single-species cultures supported with the same carbohydrates. The data were grouped based on the direction of change: up (A, C) or down (B, D).

For *S. gordonii* DL1, co-cultivation with *S. mutans* on glucose had very little impact on its transcriptome, with only 3 tRNA genes showing increased transcription and an alcohol-acetaldehyde dehydrogenase being expressed at lower levels (Table S5).

##### Co-culture on GlcN

Relative to co-cultures on glucose, co-cultivation on GlcN affected both bacterial transcriptomes in substantial ways, especially for *S. gordonii*. A total of 34 genes in *S. mutans* showed higher expression compared to *S. mutans* growing alone on GlcN, including 5 from the *slo* operon, and 6 from the TnSmu2 genomic island that encodes for enzymes required for biosynthesis of mutanobactin, a pigment important for oxidative stress resistance and for cross-kingdom interactions with *Candida albicans* (32, 33) (Fig. 6 and Table S6). Down-regulated genes (a total of 70) in *S. mutans* included *nlmC* and a variety of pathways involved in carbohydrate metabolism, including two fructose-PTS operons (SMU.113-SMU.116 and SMU.1956c-SMU.1961c), the PDH operon (SMU.1421-SMU.1425), *pflC*, and the pyruvate transporter encoded by *lrgAB*. Also down-regulated was a putative amino acid transport cluster, SMU. 930c-SMU.936. The downshift in pyruvate metabolic enzymes could be an indication that co-cultivation with *S. gordonii* reduced the availability of extracellular pyruvate for *S. mutans*, as pyruvate is a known H_2_O_2_ scavenger. The effect of H_2_O_2_ could also have impacted mutacin production as a major sensor kinase VicK, which is upregulated under oxidative stress (34), has been shown to be required for optimal mutacin gene expression in *S. mutans* (35). Compared to co-cultures grown on glucose, further reduction in all known mutacin genes was noted in co-cultures grown on GlcN (Fig. 4A and Fig. S4). Again we believe this was in part due to the effect of challisin, as higher expression of the *sgc* gene was detected in co-cultures prepared with GlcN, compared to the other two sugars. However, such increased expression in *sgc* itself is likely independent of carbohydrates, as in single-culture format *sgc* showed reduced expression on GlcN when compared to the other two carbohydrates (Fig. S4).

To our surprise, when we examined the transcripts of some of these genes via RT-qPCR by recreating the mixed-species cultures, it was clear that pyruvate metabolic genes, *lrgA* and *lytS*, SMU.1424, *pfl*, and SMU.127, were increased by co-cultivating with *S. gordonii* (Fig. 4A). The mutanobactin synthetic gene SMU.1346 and SMU.930c from the putative amino acid transporter cluster also showed increased transcript levels. Similar discrepancy between RNA-Seq and RT-qPCR was noted for certain genes in *S. gordonii* tested under the same conditions. There was an important difference between the co-cultures used in RNA-Seq and those used for RT-qPCR: the former were back-diluted 50-fold before being cultivated to mid-exponential phase for harvesting, whereas the latter were created by mixing two exponential-phase cultures in equal proportions and incubating for only 30 min (see Materials and Methods for detail). The exact reason for the difference in outcome is still unclear, however we speculate that growing together for an extended period of time (about 6-10 h, as was done with the RNA-Seq co-cultures) created markedly different environmental conditions and allowed for adaptations that would not have been experienced during the transient 30 min interaction used for the RT-qPCR experiments. These results perhaps illustrate the dynamically temporal nature of gene regulation during bacterial interactions and requires further study for clarification.

In *S. gordonii*, 170 genes were up-regulated in co-cultures with *S. mutans* growing on GlcN, compared to *S. gordonii* grown on GlcN alone. The most prominent part of the transcriptome affected by the presence of *S. mutans* belonged to the protein biosynthetic network, including 34 ribosomal proteins, 16 tRNAs, 7 amino acid-tRNA synthetases, and a number of amino acid transporter clusters (Fig. 6 and Table S6). Conversely, among the 107 genes showing reduced expression, a significant portion were involved with carbohydrate metabolism, including 4 PTS operons, the lactose/galactose cluster encoding for the tagatose pathway, the fructanase gene *fruA*, β-galactosidase *bgaA*, the glycogen operon, and others (Table S6). Contrasting this with the impact of GlcN on *S. gordonii* growing alone (Table S4), it can be inferred that in co-cultures with *S. mutans, S. gordonii* switched central metabolism from energy production into biosynthesis and growth (Fig. S5 for KEGG map). The basis for this behavior is not clear, but understanding the triggers for activation of growth by the commensal in the presence of *S. mutans* could provide some novel insights into ways to favorably modify oral biofilm ecology.

##### Co-culture on GlcNAc

Co-cultivation of these two bacteria in TV-GlcNAc resulted in a limited impact on gene expression, especially for *S. gordonii;* a result (Fig. 6 and Table S7) that was more similar to that seen with TV-glucose than TV-GlcN. For *S. mutans*, co-cultivation elevated the expression of the *slo* operon and *mntH* gene, which we again attribute at least in part to the generation of H_2_O_2_ by DL1. Again, the *nlmAB*, *nlmD* and the *nlmC*/*cipB* cluster showed significant reductions in mRNA levels. *S. gordonii* showed only 6 genes with increased expression, including a putative Mn transporter cluster. Only 4 genes showed reduced expression in *S. gordonii*, all of which were related to carbohydrate metabolism.

As a complement to the RNA-Seq analysis on mixed-species cultures, we carried out a plate-based competition assay by inoculating *S. mutans* and *S. gordonii* in close proximity, either sequentially or simultaneously, on TV base agar plates supplemented with glucose, GlcN, or GlcNAc. After incubation in the same aerobic environment as that used in the RNA-Seq study, it was clear that while these two bacteria generally co-existed on plates containing glucose or GlcNAc, they exhibited significant antagonism in the presence of GlcN (Fig. 7A). This was especially clear when one of the species was introduced 1 day earlier than the other, where the earlier colonizer consistently outcompeted the other species. These observations are consistent with the drastic shifts in transcriptomes shown by both bacteria when co-cultivated on GlcN.

**Fig. 7.**
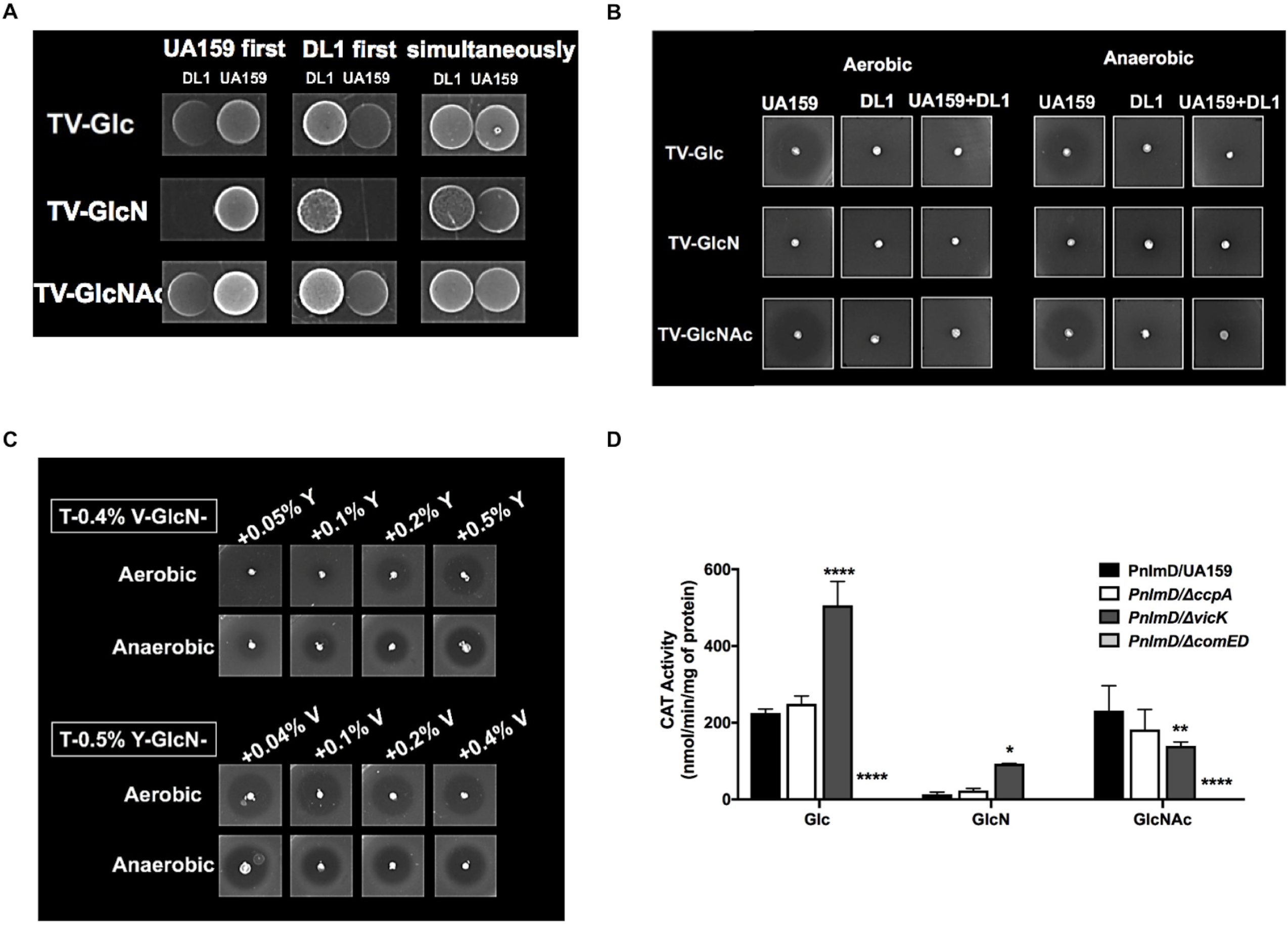
Bacteriocin expression by *S. mutans* is regulated by multiple environmental factors. (A) *S. mutans* UA159 and *S. gordonii* DL1 were spotted on TV-agar plates supported by glucose, GlcN, or GlcNAc, either simultaneously or 1 day apart, followed by incubation in an aerobic environment supplemented with 5% CO_2_. (B, C) UA159 was tested in a deferred antagonism assay under specified nutritional and atmosphereic conditions, in the presence (B) or absence (B, C) of DL1. (D) The expression of a P*nlmD∷cat* fusion was studied in the wild-type UA159 and mutant derivatives deficient in *ccpA*, *vicK*, or *comED*, with carbohydrate being the other variable. Each bar represents the average of three biological repeats, with error bar denoting standard deviation. Asterisks indicate statistical significance in comparisons to the wild type growing on the same carbohydrates according to two-way ANOVA (* *p* <0.05, ** *p* <0.01, *** *p* <0.001, **** *p* <0.0001).

### The effects of amino sugars on bacteriocin production is impacted by nitrogen source and redox signal

As noted earlier, growth on TV-GlcN resulted in reduced expression of mutacin gene clusters in *S. mutans* (Fig. 4A and Table S1), and this contradicted our previous findings on TY agar plates, where amino sugars appeared to increase the ability of *S. mutans* to kill in a deferred antagonism assay (13). First, we carried out a deferred antagonism assay on TV agar plates containing glucose, GlcN or GlcNAc using *S. sanguinis* SK150 as an indicator strain, as in (13). The results (Fig. 7B) showed substantially less killing by mutacins of *S. mutans* only on plates containing GlcN, in aerobic or anaerobic environments, confirming the results observed by RNA-Seq analysis. The co-inoculation of *S. gordonii* together with *S. mutans* suppressed the production of mutacins under all conditions, which is also consistent with the results of RNA-Seq showing reduced expression of *nlm* genes in co-cultures.

The main difference between TV- and TY-base media is the lack of yeast extract in the former and the lack of a vitamin mixture in the latter. To determine which factor was responsible for the differences in bacteriocin production, we created a modified TV agar by adding increasing amounts of yeast extract (0.05%-0.5%) to the medium, and a modified TY agar with increasing amounts of vitamin mix (0.04%-0.4%); all with the same amounts of GlcN as the sole carbohydrate. When we repeated the deferred antagonism assay on these plates, *S. mutans* (Fig. 7C) showed a concentration-dependent response to yeast extract in the size of the zones of inhibition formed on TV media, producing greater inhibition zones with higher concentrations of yeast extract. Conversely, addition of increasing amounts of the vitamin mix to TY agar had no impact on antagonism. Previous research (36–38) has suggested that production of mutacins and the expression of relevant genes require both nitrogen-rich ingredients, such as yeast extract, and specific carbohydrates, each provided at optimal levels. Here, we show that amino sugars can impact mutacin production by *S. mutans* in both directions, dependent on the overall nutritional state of the bacterium.

To further explore the transcriptional regulation of mutacins by carbohydrates, we utilized a P*nlmD*∷*cat* reporter fusion (39) and studied the response of the *nlmD* promoter to amino sugars. Based on behaviors seen in the RNA-Seq experiments, we also included mutant strains lacking CcpA (40), VicK (35), or the CSP-responsive two-component system ComED required for mutacin expression (41). The study was carried out using TV base medium in an aerobic atmosphere containing 5% CO_2_. The results of the CAT assays in the wild-type UA159 (Fig. 7D) showed a clear reduction in *nlmD* expression in GlcN-grown cultures, relative to glucose cultures, whereas little difference was noted when using GlcNAc. Elimination of catabolite control protein CcpA had no impact on *nlmD* expression on any of the carbohydrates tested. However, loss of the histidine kinase VicK altered expression of *nlmD* in all three conditions: the promoter activity more than doubled with glucose or GlcN, yet it showed significant reductions with GlcNAc. As a control, loss of the ComED two-component system caused complete loss of *nlmD* promoter activity in all conditions. Mutacins are regulated by various environmental factors, and by multiple genetic systems that often interact and overlap in functionality (38, 42). Previous research has shown the capacity for the VicRK system to affect the expression and function of the ComED system (35), as well as effects of VicRK on oxidative stress (34) and in trans-phosphorylating GcrR/CovR in a Mn-dependent manner (42); GcrR/CovR is an orphan response regulator involved in acid tolerance response and biofilm development (43, 44). Together, these results and the RNA-Seq analysis suggest that the intersection of redox signaling, essential ions, and nutritional status likely drives the bacterium to balance the needs for growth, energy production, and defense.

### Concluding remarks

Release of pyruvate during carbohydrate metabolism has been reported in a few oral bacteria, including *S. mutans* and commensal bacteria *S. gordonii* and *S. sanguinis*. Overflow metabolism allows the bacterium to achieve rapid growth and lactic acid production when carbon sources are plentiful, and subsequent uptake of extracellular pyruvate allows extended generation of ATP when resources become limited. Since *S. gordonii* also releases H_2_O_2_ via the action of SpxB on pyruvate, and pyruvate is a H_2_O_2_ scavenger, the role of extracellular pyruvate in the physiology and ecology of commensal streptococci warrants further study. However, one reasonable explanation is that release of H_2_O_2_ evolved as a weapon against bacteria such as *S. mutans*, while pyruvate serves to protect the commensal from self-harm (25). In the case of *S. mutans*, which lacks SpxB and does not produce significant quantities of H_2_O_2_, pyruvate should also provide some protection from oxidative damage from H_2_O_2_. Utilizing RNA-Seq instead of a reductive approach, we were able to identify the impact of two amino sugars on pyruvate metabolism on *S. mutans* and *S. gordonii*, individually and when growing together. Through release of metabolic byproducts, namely ammonia and acetate, amino sugars could initiate significant perturbations in pyruvate metabolism, a homeostasis that is subjected to influence by environmental factors, including carbohydrate and nitrogen sources, oxidative stressors (O_2_, H_2_O_2_), and essential metal ions. These adaptations in turn result in carbon flux redistribution among energy and acid production, growth, and biosynthetic pathways. While we continue improving this working model, these findings allow us to expand our insights into amino sugar metabolism by other species of the dental biofilm, and explore the potential of utilizing metabolic modulation towards caries management and intervention.

## MATERIALS AND METHODS

### Bacterial strains and culture conditions

Three bacterial species were used in this study: *S. mutans* UA159 and derivatives therefrom, *S. gordonii* DL1, and *S. sanguinis* (strain SK150). Brain heart infusion (BHI, Difco Laboratories, Detroit, MI) medium was used in both liquid form and agar for maintenance of bacterial strains. For experiments carried out with carbohydrates as the sole variable, a semi-defined Tryptone-Vitamin (TV) based medium (45), a TY base medium (3% Tryptone, 0.5% yeast extract), or a modified chemically defined FMC medium (17) was used with addition of the specified amounts of carbohydrates. TV and TY base media could be prepared in advance, with carbohydrates added immediately before use and the pH of the medium adjusted to 7.0 after addition of the carbohydrate. All bacterial cultures were maintained at 37°C in an aerobic atmosphere supplemented with 5% CO_2_, or an anaerobic environment established using an AnaeroPack (Thermo Fisher Scientific, Waltham, MA).

For RNA extraction, and for most assays performed in this report, bacterial strains UA159 and DL1 were first cultivated overnight, each with at least 3 individual cultures, on a TV medium supplemented with 20 mM glucose, GlcN, or GlcNAc, then diluted 1:20 ratio into fresh TV medium containing the same carbohydrate as was used to grow the inocula. When the optical density (OD_600_) of an individual culture reached 0.5, a portion of the culture was placed on ice temporarily before use. The rest of the culture was harvested by centrifugation, treated with the RNAprotect Bacteria Reagent (Qiagen, Germantown, MD) and stored at −80°C. Once all cultures were ready on ice, UA159 and DL1 that had been grown on the same carbohydrates were mixed in a 20:1 ratio based on their OD_600_ values, then the mixture was diluted 1:50 ratio into the same medium used to prepare the inocula. To ensure that approximately equal numbers of bacteria from each species were present at the end of the cultivation, the ratio at which these two strains were being mixed was determined empirically by enumerating their final CFU on agar plates with and without bacitracin. After the dual-species culture reached OD_600_ = 0.5, cells were harvested as before and stored at −80°C. For confirmation in differential gene expression after RNA-Seq analysis, the dual-species cultures were recreated as before, with one exception: once the single-species cultures reached OD_600_ = 0.5, the two bacteria were mixed in a 1:1 ratio, without dilution, and incubated for 30 min before harvesting for RNA extraction. We have noted substantial day-to-day variability in the proportions of these two bacteria in co-cultures harvested for RNA-Seq, this new approach allowed us a better control in the amounts of bacteria engaged in interactions. This small change in culture method however resulted in significant differences in expression of a number of genes by both bacteria.

### RNA extraction, RNA-Seq, data analysis, and RT-qPCR

Total RNA samples from single- and dual-species cultures were extracted using the RNeasy mini kit (Qiagen), by following an established protocol detailed elsewhere (16). To enrich mRNA by removing 16S and 23S rRNAs from each RNA sample, 10 μg of total RNA was treated, twice, using the MICROBExpress bacterial mRNA enrichment kit (Ambion Life Technologies, Grand Island, NY), as directed by the suppliers. The quality of enriched mRNA samples was assessed using an Agilent Bioanalyzer (Agilent Technologies, Santa Clara, CA). cDNA libraries were constructed from 100 ng of enriched mRNA samples by using an NEBNext Ultra directional RNA library prep kit for Illumina, and an NEBNext multiplex oligonucleotides for Illumina (New England BioLabs, Ipswich, MA) for barcoding. Parallel deep sequencing was carried out by the NextGen DNA Sequencing Core Laboratory of ICBR at the University of Florida (Gainesville, FL). The high-throughput sequencing data from this study have been deposited with Gene Expression Omnibus (GEO) and assigned accession number GSE152021.

The analysis of the RNA-Seq expression data was conducted on R (46) versions 3.3.2 “Sincere Pumpkin” and 3.3.3 “Another Canoe”. The compositional matrix of the expression data was normalized with the “voom” (47) function from the R package “limma” (48). These normalized datasets were used to generate heatmaps, correlation plots, and PCA plots. Correlation and heatmaps of the compositional matrix were plotted with the “corrplot” package (49) and the enhanced “heatmap.2” function from the “gplots” (50) package, respectively. The data for PCA plots were extracted from the “prcomp” base function output and used to generate plots with the package “ggplot2” (51) in combination with the “ellipse” (52) function to produce the ellipse confidence regions. The overlapping set of genes within different conditions were plotted as Venn diagrams with the function “vennPlot” from the package “systemPipeR” (53). The statistical analysis of the expression data was conducted following the DESeq2 pipeline [4] from the homonymous R package. In order to follow pairwise comparisons required by the experiment, the contrasts were parsed to the pipeline by the results function. The pipeline output data included both a *p*-value and a multiple comparison adjusted *p*-value. A false discovery rate (FDR) of 0.01 was used as cutoff value to call the significant genes in the results, and the volcano plots as well. Subsequently, KEGG functional pathways and gene ontology (GO) terms were generated individually for all datasets using DAVID bioinformatic resources (54). Pathway maps were generated using the KEGG Mapping tools from Kyoto Encyclopedia of Genes and Genomes (KEGG) resources (55).

Reverse transcription quantitative PCR (RT-qPCR) was carried out for confirmation of selected genes after RNA-Seq analysis. RT was carried out using the iScript cDNA synthesis kit (Bio-Rad, Hercules, CA) on 0.5 μg of total RNA in a 10-μL reaction, with each gene-specific, anti-sense primer added at 200 nM (Table S8). qPCR was performed on a CFX96 Touch Real-Time PCR Detection System, using the SsoAdvanced Universal SYBR Green Supermix (Bio-Rad) and a 1:10 dilution of the original RT reaction as template for amplification. cDNA levels were calculated using the ΔΔCq method, with a housekeeping gene *gyrA* from each bacterium as the internal control. Considering the relatively close phylogenetic relationship of *S. mutans* and *S. gordonii*, additional control experiments were performed to assess the stringency of RT-qPCR reactions when using RNA samples from mixed-species co-cultures. Briefly, anti-sense primers for a select number of *S. mutans* genes were used in RT reactions together with RNA extracted from 3 *S. gordonii* cultures (TV-Glc), while in 3 separate reactions, primers for *S. gordonii* genes were used to prime RNA from *S. mutans* cultures (TV-Glc). qPCR results derived from these RT reactions confirmed the absence of any significant cross-species priming in reactions that used RNA samples from mixed-species cultures, with the majority of the Cq values being >30 under the current experimental setup (data not shown).

### Deferred antagonism assay

Overnight cultures of *S. mutans* UA159 and *S. gordonii* DL1 grown in TV containing 20 mM Glc, GlcN or GlcNAc were each diluted into fresh TV medium containing the same carbohydrate, and incubated until OD_600_ reached 0.5. UA159 alone, DL1 alone, or UA159 together with DL1 at a ratio of 1:1 was inoculated by stabbing onto TV agar plates prepared with 20 mM Glc, GlcN or GlcNAc, in accordance with carbohydrates used in cultures. After 24 h of incubation at 37°C, each plate was overlaid with 8 mL of warm soft agar (0.75% TV agar, prepared with the same carbohydrate as was in the plate) containing 10^7^ CFU of *S. sanguinis* SK150 as an indicator strain. Plates were incubated for an additional 24 h before measuring zones of inhibition.

To understand the different impact of TV and TY media on production of mutacins by *S. mutans*, various amounts of yeast extract, ranging from 0.05% to 0.5%, were added into TV base medium containing 20 mM of GlcN. For comparison, increasing amounts (0.04% to 0.4%) of the stock TV vitamin mix (10 μg/mL *p*-aminobenzoic acid, 50 μg/mL thiamine-HCl, 250 μg/mL nicotinamide, and 50 μg/mL riboflavin) were added into TY base medium supplemented with 20 mM of GlcN. Using agar plates prepared with these modified TV and TY media, a UA159 culture from the exponential phase was used as above in a deferred antagonism assay with *S. sanguinis* SK150.

### Plate-based competition assay

TV-based cultures of UA159 and DL1 from the exponential phase were prepared as above, using 3 different carbohydrates. Six μL each of the cell suspensions of UA159 and DL1, prepared with the same carbohydrate, were spotted in close proximity on TV agar plates formulated with 20 mM Glc, GlcN or GlcNAc, in accordance with the carbohydrates used in the cultures. Depending on the experimental design, UA159 or DL1 was inoculated alone first, incubated for 24 h, followed by the other strain; or they were spotted simultaneously. Plates were incubated in aerobic conditions with 5% CO_2_ for another 24 h at 37°C before being photographed.

### Promoter∷reporter fusion and enzymatic assays

Chloramphenicol acetyltransferase (CAT) assays were carried out to quantify promoter activities according to an established protocol (56). *S. mutans* derivatives containing an *nlmD* promoter∷*cat* fusion (39) were cultivated to mid-exponential phase (OD_600_ = 0.5) in TV prepared with 20 mM Glc, GlcN or GlcNAc, before being harvested for assays. To assess the expression of a P*lrgA*∷*gfp* (green fluorescent protein) reporter fusion, the *S. mutans* derivative hosting the fusion plasmid (18) was inoculated into a synthetic FMC medium supported by 10 mM Glc, GlcN or GlcNAc, and monitored for both optical density and green fluorescence signals for 30 h in a fluorescence-capable plate reader (57).

To measure pyruvate levels in supernatant fluids, overnight BHI cultures of *S. mutans* were washed and resuspended in FMC base medium, followed by 1:20 dilution into a FMC constituted with 10 mM Glc or GlcN. To avoid the long lag often associated with transferring UA159 from glucose-based media onto FMC-GlcNAc (14), we prepared a set of TV-GlcNAc cultures as starters and diluted them similarly into FMC containing 10 mM GlcNAc. At various time points (~1-h interval), aliquots of the bacterial culture were removed for measurement of OD_600_, or spun down to harvest the supernates. Pyruvate levels in these culture supernates were measured using an EnzyChrom pyruvate assay kit (BioAssay Systems, Hayward, CA) by following the supplier’s instructions. The results were plotted together with their corresponding OD_600_ values. FMC was chosen over TV in pyruvate-related experiments to minimize media components interfering with the fluorescence signal and the pyruvate assay.

## ACKNOWLEDGEMENTS

This study was supported by DE12236 and DE25832 from NIDCR. We thank Dr. Sang-Joon Ahn for useful discussions and providing the GFP reporter strain.

